# Systematic Review on Barriers and Facilitators for Access to Diabetic Retinopathy Screening Services

**DOI:** 10.1101/335638

**Authors:** M.M. Prabhath Nishantha Piyasena, Gudlavalleti Venkata S. Murthy, Jennifer L.Y. Yip, Clare Gilbert, Maria Zuurmond, Tunde Peto, Iris Gordon, Suwin Hewage, Sureshkumar Kamalakannan

## Abstract

**Objectives:** The aim of this systematic review is to identify the barriers/enablers for the people with diabetes (PwDM) in accessing DRS services (DRSS) and challenges/facilitators for the providers.

**Background:** Diabetic retinopathy (DR) can lead to visual impairment and blindness if not detected and treated in time. Achievement of an acceptable level of screening coverage is a challenge in any setting. Both patient-related and provider-related factors affect provision of DR screening (DRS) and uptake of services.

**Methods:** We searched MEDLINE, Embase, CENTRAL in the Cochrane Library from the databases start date to September 2016. We included the studies reported on barriers and enablers to access DRS by PwDM and studies which have assessed barriers or facilitators experienced by the providers in provision of DRSS. We identified and classified the studies that used quantitative or qualitative methods for data collection and analysis in reporting themes of barriers and enablers.

**Main Results:** We included 63 studies primarily describing the barriers and enablers. The findings of these studies were based on PwDM from different socio-economic backgrounds and different levels of income settings. Most of the studies were from high income settings (48/63, 76.2%) and cross sectional in design (49/63, 77.8%). From the perspectives of users, lack of knowledge, attitude, awareness and motivation were identified as major barriers to access DRSS. The enablers to access DRSS were fear of blindness, proximity of screening facility, experiences of vision loss and being concerned of eye complications. Providers often mentioned that lack of awareness and knowledge among the PwDM was the main barrier to access. In their perspective lack of skilled human resources, training programs and infrastructure of retinal imaging and cost of services were the main obstacles in provision of screening services.

**Conclusion:** Knowing the barriers to access DRS is a pre-requisite in development of a successful screening program. The awareness, knowledge and attitude of the consumers, availability of skilled human resources and infrastructure emerged as the major barriers to access to DRS in any income setting.

## SECTION 1

### Introduction

Diabetes mellitus (DM) is one of the most prevalent non-communicable diseases which imposes a significant impact on health systems. The International Diabetes Federation (IDF) estimated that there were 425 million people with diabetes (PwDM) in the world and this will increases to 629 million by 2045.(1) It has been emphasised that efforts should be made to prevent the complications of DM as per the targets set in St Vincent declaration in 1989.(2) Diabetic retinopathy (DR) is a common microvascular complication caused by chronic hyperglycaemia. Blindness due to DR is common among the working age populations and it is becoming a global issue due to rising prevalence of DM.(3) Though proportion is low, expenditure related to DR is a burden to any health system.(4)

Optimal access to health care was defined by Rogers et al in 1999 as *“providing the right service at the right time in the right place”.*(5) Access to health care remained a vague concept, till recently, impeding the work of health care policy makers. In generic literature it is mentioned that access has multi-dimensions and it is not merely the entry in to the healthcare system.(6) Further it is an outcome of people’s potential to use health care and manifestations of patients actual use.(7) Donabedian has observed that the proof of access is use of service, not simply the presence of a facility.(8) Some authors argue that it depends on acceptability of the services as well.(9,10) Penchansky and Thomas described the concept of access as the *“degree of fit”* between clients and the health system.(6) It is mentioned that inequalities have been observed in detection and treatment of DR which require multi-sectoral engagement.(11) Further universal coverage cannot be achieved without addressing the barriers.(12) One review mentioned that there are many reasons for underutilization of eyecare and that the risk of blindness varies with the context.(13) It has been shown that culturally competent care should be delivered in a diverse patient community overcoming the sociocultural barriers.(14,15)

Screening of DR can be done opportunistically or proactively. Current literature shows that proper disease control, DRS and early identification and treatment of pathologies will reduce progression of sight threatening DR (STDR).(16–20) Awareness of detecting DR at a symptomless stage is a key factor in uptake and regular follow up of DRS services (DRSS). There are many obstacles for implementation and maintaining satisfactory level of uptake in DRS in program level. One important approach to address this issue is identification of challenges and incentives in the system in advance. Defining a barrier will enable implementation of public health strategies to improve access.(6) A barrier could lead to a different outcome for a certain community such as barriers for people with disabilities.(21) In system assessments authors mentioned that economic and logistic reasons hinder the provision of screening services.(22) It is mentioned that effective strategies are underutilised in developing countries to overcome barriers.(23) Therefore it is necessary to understand the potential barriers in accessing and challenges in provision of DRSS in any health system.

#### Conceptual framework

The successful uptake of DRSS depends on the personal factors related to the consumer as an agency. These factors may be modifiable or not modifiable according to the environment. The required behavioural change techniques for a target population could be hypothesised using various behavioural models. The “*social cognitive theory*” explains how persons acquire and maintain specific behavioural patterns and it provides the basis of most intervention strategies to overcome a defined barrier.(24) A person’s behaviour influences and is influenced by personal factors and the social environment (‘Reciprocal determinism’) (25) (S1 Fig 1). This will lead to self-efficacy of the person to achieve confidence for performing a particular behaviour. Therefore, assessment of behavioural patterns and perceived barriers in accessing DRSS may be useful in developing a successful DRS program (DRSP) in any context.

There are no published reviews related to this topic. One protocol was available however it is yet to be published.(26) Most of the individual studies had provided the evidence of barriers to access DRSS according to the typology of barriers. The processes related to DRS uptake can be considered in three levels i.e., service user, service provider and eyecare system. Therefore, in this review we categorised the reported themes or variables under above categories. This review was specifically assessed the barriers to access DRSS at established health care facilities and challenges / barriers faced by the providers in those institutions. In broad definitions, barriers to access to DRS is not only limited to the access issues at the point of delivery, but it also involves all the steps which take place starting from perceptions of a PwDM at one end to the whole eye care system at the other which are inter-related and connected to each other.

#### Objectives

The overall aim of the review was to explore barriers to access DRS. The review has the following specific objectives.

-To assess the barriers and enablers to uptake of DRSS by PwDM.
-To assess the challenges faced by the services providers in provision of DRSS and to identify the facilitators for development of a DRSP.

The secondary objectives of this review were;

-To assess the socio-economic factors that could affect DRS uptake.
-To assess the barriers or enablers to develop DRSP in a health care system.

## SECTION 2

### Methods

We included studies that focused on assessing barriers / challenges and enablers / incentives to access DRS. In addition, we found studies that described factors affecting the uptake of DRSS. Following criteria were used for assessment of eligibility of the studies. (There is no protocol registration for this review and PRISMA checklist was included as S1 Table 1).

Inclusion of studies -

-Consumers - The studies which have assessed the barriers at group or individual level of PwDM at or had been referred to a permanent health care facility for DRS were considered.
-Providers - study participants were providers who have direct contact with PwDM in a permanent health care institution and / or decision makers / other stake holder involved in decision making on DR.

Exclusion of studies -

-Studies which included general population as the study sample without specifying the status of DM.
-Absence of standard diagnostic criteria for DM.
-Assessment of the barriers for eye care in general without specifying DRS.
-Assessment of barriers for DM complications in general without specifying the barriers for DR.

We did not restrict the studies for inclusion by study design. We included studies that used qualitative, quantitative and mixed methods.

#### Type of participants

We included the studies that have covered PwDM who were attending a diabetic medical care or an eye care facility.

#### Type of interventions

We included studies that delivered or considered DRSS primarily at an established health care facility. We defined the DRS as performance of dilated retinal screening using imaging (digital / colour films) or by direct / indirect ophthalmoscopy by a trained / skilled eye care professional (preferably an ophthalmologist / retinologist) to identify the signs of DR.

#### Type of outcome measures

We defined access as all level of factors affecting the processes of DRS in a health care facility.

#### Phenomena of interest

We included the studies which have assessed barriers or enablers to access DRS by PwDM and challenges or incentives faced by providers in provision of screening services in current screening programs or at opportunistic screening.

#### Search method for the identification of studies

We searched Ovid MEDLINE, Embase and CENTRAL in the Cochrane Library from the databases inception up to 15^th^ September 2016. The search strategy was developed by an information specialist from Cochrane Eyes and Vision (IG) (search terms available in S2 Table 2). We did not use any filtering methods to limit the results by study design, year of publication of language. This yielded a comprehensive coverage of published articles. However due to resource restrains we were not able to translate any non-English reports.

#### Data collection and analysis

Two reviewers (PN and SK) independently assessed the eligibility of inclusion by going through titles and abstracts of 13,082 articles after, importing them to an EndNote^®^ library. The potential articles (full papers as identified by either or both reviewers) were retrieved from publishers. These papers were then assessed independently by two reviewers (PN and SK). Disagreements between the reviewers were resolved by a 3^rd^ arbitrary reviewer (GV). Reviewers assessed full papers independently to retrieve accurate data. We aimed to include all relevant studies from different income settings to avoid bias in selecting articles. Therefore, we could interpret a range of barriers themes with a greater variation and a greater conceptual diversity.

#### Data extraction and management

We developed an MS Office Excel^®^ data sheet to directly transfer extracted data from full articles. The topics to be extracted were developed according the “Strengthening the Reporting of Observational Studies in Epidemiology” statement (STROBE) and modelling has been done according to the review question.(27) The accuracy of extracted data was cross-checked by a third reviewer (SH).

We extracted information on first author’s name, year of publication, country of study (by income category), place of the study, sample size, gender distribution, mean age, method of diagnosing DM, level of DM and DR of the participants and method of DRS in the 1^st^ set of data. In the next step we collected information on type of study design, objective, study setting, data sources, sampling strategy, time period when study was conducted. The methodological quality assessment and applicability for review question were done separately as subsequently described. We collected the results and main outcomes of the study according to the review question.

In the synthesis of evidence “informants” were authors of the individual studies rather than the participants. The authors’ interpretations were presented as narrative themes supported by numerical values of statistical significance levels wherever available.

While authors’ interpretations were primarily collected from results section of each paper, author interpretations were sometimes also found in the discussions sections and these were also extracted when relevant and well supported by data.

#### Assessment of risk of bias in included articles

We carried out the risk of bias and quality assessment according to the guidelines of critical appraisal of skills program (CASP) tools for case-control, qualitative, cohort and randomised controlled study designs (28) and National Institute of Health, United States quality assessment tool for observational cohort and cross sectional study designs (NIH-QAT) for cross sectional study designs.(29) Two reviewers (SK and PN) independently applied set of quality criteria to each included study. We appraised how well the individual studies conducted which contributed to narrative synthesis using the above tools. Emphasis was given more over the applicability of the study according to the inclusion criteria. It has been noted that applicability to review question was the main concern in the synthesis rather than the overall level of quality of a study (S3 Table).

#### Assessment of methodological limitations

When several studies with varied methodological limitations contributed to a finding, we made an overall judgement about the distribution of strengths and weakness of the study rather than for individual components in the tools.

#### Assessing coherence

We assessed the coherence of each review finding by looking at extent to which we could identify a clear pattern across the data contributed by each of the individual studies. This was supported by when clarity of the themes was consistent across different contexts and the variations were explained by the study authors according to the data collected, when supported by numerical data (odds ratios). This was further strengthened when findings were drawn from different settings

#### Data synthesis

Most of the eligible studies were observational and descriptive in nature hence narrative reporting approach was used in this review. We analysed and synthesised the descriptive and qualitative data narratively supported by other associated variables with levels of statistical significance. We described the barriers and enablers according to the dimensions of the typology of barriers and this was tied with processes involved in DRS. When describing the themes, we did not re-phrase the findings or conclusions mentioned by authors. We used imputations up to a certain degree in describing barrier themes.

Considering the participants of the studies, the themes that emerged were divided in to three categories complying with the objectives of the systematic reviews. Those are consumer perspectives, provider perspectives and system factors.

### RESULTS

#### Results of the search

Search and study selection procedures are summarized in the PRISMA flow diagram (Fig 1). The search identified a total of 13,082 records. Duplicate records were removed, and we assessed 13,055 reports for potential inclusion in the review. We excluded 12,954 records based on the information given in the title and abstract. After assessing the full-text of 101 reports of studies, we excluded 38 studies which did not meet the inclusion criteria and included a total of 63 studies in the review.

**Fig 1.**
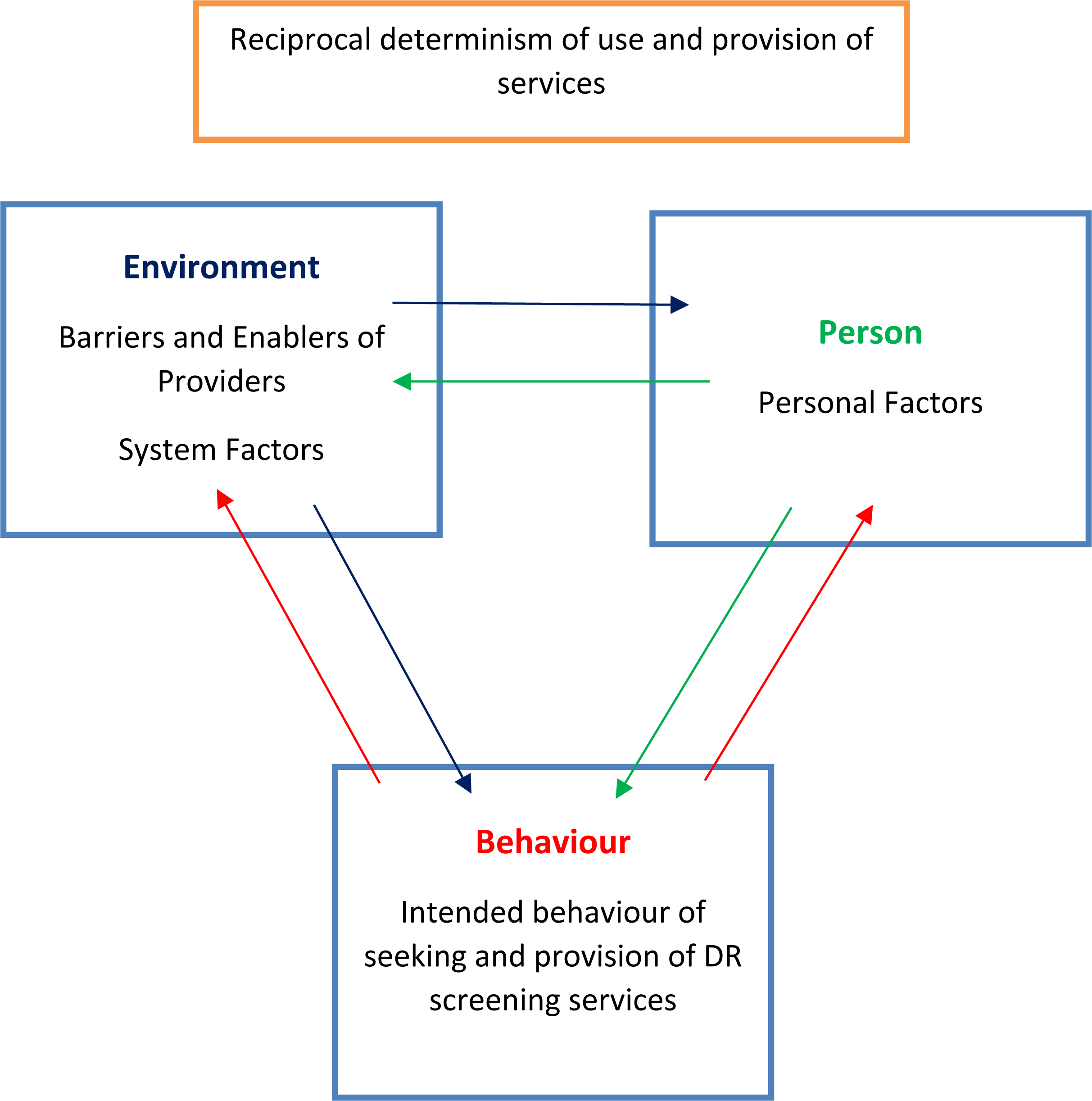
Schematic diagram of “Social Cognitive Theory”

#### Methodological quality of the studies

The methodological quality assessments of included studies were presented in S3 Tables 1 to 5 according to the study design. In the included cross-sectional studies, 98% (49/50) of the studies clearly stated study objective matching the review question. Sample size justification was not available in 52% (26/50) of the studies. Participation of eligible persons at least 50% was not seen in seven (7/50, 14%) studies. Four of the studies (4/50, 8%) have not recruited the participants from a similar population. The outcome measures were not clearly defined in 14 studies (14/50, 28%) and confounders were not adjusted in ten studies (10/50, 20%).

Acceptable method of recruitment of the cohort was not followed in all three included cohort studies. In included randomised controlled study designs, applicability of the results to the PwDM was not observed in two studies (2/3, 66%). In qualitative study designs, most of the quality assessment criteria were met except, relationship between researcher and the participants were not adequately considered in two studies (2/5, 40%).

#### Overview of the included studies

We identified a total of 13082 titles and abstracts and considered 101 full text papers for inclusion in this review. Sixty-three (63/101, 62.3%) studies were eligible for inclusion in the narrative review according to the objectives. The S4 Table file contains the details of participants and settings.

#### Included studies

All the studies main group of respondents were PwDM. Some authors have sought barrier perspectives form providers as well. Forty-eight studies (76.2%, 48/63) described barriers related to consumers, providers and eye care system, 3 studies on consumers and system (4.8%, 3/63), 1 study on provider and system (1.6%, 1/63) and 9 studies on consumer and provider (14.3%, 9/63). Only 2 studies (3.2%, 2/63) described barriers of consumers only and no study has focused only on providers. Two studies reported the outcome as a review.(30)(31).

#### Settings-by income

Only three (4.8%, 3/63) (30,32,33) studies were from low income (LI African countries, Tanzania, Nepal) countries (LIC). Seven were from lower middle (11.1%, 7/63) (Indonesia, India, Yemen, Kenya, Myanmar, Nigeria)(LMIC) (34–40), 5 upper middle (7.9%, 5/63) (Turkey, Iran, Mediterranean countries, China)(UMIC) (41–45) and 48 high income (76.2%, 48/63) settings (HIC)(16/63-25.4% from United Kingdom, 19/63-30.2% from United States, Other-Germany, France, Ireland, Singapore, Canada, Oman, Hong Kong, South Korea, Australia, Taiwan, Italy and Netherland). (46–93).

#### Setting-by type of institution

Most of the data collections were done under the primary level general practices, local clinics and primary care clinics (15/63, 23.8%).(38,52,53,56,57,60,63,65,66,68,81,85,87,88,90). There were 12 population-based studies (12/63, 19.0%). (39,45,48,49,58,59,72,73,77,84,86,93) Eight studies were conducted at tertiary level institutions (8/63, 12.7%: 7 eye clinics and 1 endocrinology clinic).(32– 34,40,41,44,54,55) Five studies were conducted in existing DR screening programs (5/63, 7.9%).(61,82,83,91,92)

Eight studies were conducted at secondary level medical and diabetes clinics (8/63, 12.7%). (37,42,47,51,64,67,71,80) Five studies were conducted analysing existing data bases (5/63, 7.9%). (69,74–76,79) There was one study where authors did not mention about the setting (70) and two studies reported the barriers as a review.(30,31) Two studies collected the sample of PwDM at a screening camp and at an annual campaign.(35,50) One study was conducted in a an ambulatory clinic based at a nursing home.(46) Three studies conducted at eye clinics (3/63, 4.8%; 2 at general eye clinic (36,89) and 1 optometry practice (78)). One study conducted using at a model of not for profit health model. (62)

#### Included studies

This analysis mainly comprised of cross sectional observational studies. In the included 63 studies, there were 49 (49/63, 77.8%) cross sectional observational studies (observational 25, retrospective studies 8, postal surveys 1, telephone interview 2, 1 mixed method audit and 12 population-based studies (12/63, 19.0%), Other study designs were 3 controlled trials (3/63, 4.8%),1 case control study (1/63, 1.5%), 3 cohort studies (3/63, 4.7%) and 5 qualitative studies (5/63, 7.9%). There were 2 reviews (2/63, 3.2%) in the included studies.

#### Synthesis

Our main objective was to identify barriers or facilitators to access DRS. Our findings are summarised in the S5 Tables files according to the country income.

#### Narrative summary - Barriers

The following main themes were derived from descriptive and qualitative studies (S5 Table 1 to 4).

#### LIC

The most prominent barrier to access DRS among the consumers in LIC were lack of knowledge on DM eye complications, lack of awareness about importance of eye examination and lack of knowledge about availability of eye clinics. Among providers, main challenges were lack of skilled human resources and lack of access to DR imaging and treatment infrastructure. Further non-existence of a referral system and lack of multi-disciplinary care approach were barriers to provision of DRSS. In LIC lack of a national policy and competing disease priority environments were the main obstacles in the system (S5 Table 1).

#### LMIC

Consumers’ barriers related to knowledge and awareness could be observed in the LMIC as well. Additionally, financial barriers and disabilities emerged as themes of barriers. In providers perspectives lack of DM health education and financial barriers were the main barriers. Lack of human resources, uneven distribution of skilled personnel, lack of availability of equipment and treatment facilities, low referral rates and time constrains in busy eye clinics were the main challenges faced by providers in LMICs in provision of DRSS. The information given by the provider to PwDM and training of non-ophthalmologist human resources were the main incentives mentioned by the providers. In system analysis lack of training, accessible eye centres and lack of epidemiological studies were emerged as main barriers (S5 Table 2).

#### UMIC

The lack of awareness and knowledge on DR emerged as the main barrier among the PwDM in UMIC. Poor physician-patient communication was also a barrier in these countries. In provider perspectives scarce human resources, lack of training, high number of PwDM were the main challenges faced. In the system analysis limitations in prevention and health promotion, civil unrest, disparity in urban and rural services, lack of transportation and problems in insurance schemes were the main barriers to accessing DRSS (S5 Table 3).

#### HIC

In HIC living alone, problems in mobility, problems in accessing general practitioner, effects of mydriasis prohibiting driving, reluctance to change behaviour, disliking the method of examination, change of residence, problems in securing appointments, being employed, extended vacations were observed as the main barriers among the consumers. In some HIC, lack of knowledge regarding eye examination, lack of awareness of eye care, lack of flexibility in adjusting attitude and behaviour and lack of understanding of rationale and importance of annual eye examination were observed as barriers to access DRSS. Even in the HIC socio-economic inequalities, poor communication skills, social deprivation and poorer literacy were barriers to access DRSS among some communities.

In the providers perspectives; level of the experience of the screener, lack of attention by the general practitioners, non-adherence to guidelines, lack of information provided to patients, lack of physician recommendations, limited knowledge on DR among the health professionals, long waiting time (large number of patients per doctor), failure to refer by general practitioner, perceptions of side effects of mydriasis, limited knowledge-attitude and practice of physicians, limited experience in using ophthalmoscope, long waiting time for treatment, lack of communication between screening services and practices were mentioned as barriers. Providers mentioned that problems associated with consumers such as confused and immobile patients, unawareness of importance of mydriasis, poor physician-patient communication, different perceptions in making appointments, after effects of mydriasis, fear of laser, wrong assumption on patient’s level of knowledge could hinder to access DRSS.

In HIC system analysis lack of understanding among the specialities, frequent change of staff, lack of human resources, unavailability of medical records, lack of adequately trained optometrists, lack of proper referral and reminding system, lack of insurance coverage, financial barriers, unavailability of national programs, problems in transportation were barriers to access or provision of DRSS (S5 Table 4).

#### Narrative summary - Enablers/Incentives

Themes of enablers / incentives are summarised in the S5 Tables 1 to 4 files according to the country income.

#### LIC

In LIC settings users’ knowledge on DR, having a family member with DM and prior fundus examination were incentives to attend DRS. Provision of imaging and treatment infrastructure, increased human resources, provision of training on retinal care and prioritisation of development of subspecialties were mentioned as enablers for the providers (S5 Table 1).

#### LMIC

The incentives to uptake of services by the users were presence of symptoms, more severe DM and comorbidities, better understanding of risk factors, patient satisfaction over the modality of screening and presence of visual impairment / blindness. Training of non-ophthalmologist physicians on DRS, availability of fundus camera, educational strategies aimed both patients and physicians, a reminder of the serious consequences of the failure to undergo DRS and public health education using media were emerged as incentives to improve uptake of DRSS in LMIC (S5 Table2).

#### UMIC

Higher literacy, person’s concern about the vision loss, severe DR stage and having knowledge on DR were the main facilitators for users in DRSS uptake. In UMIC awareness among the physicians DM complications, availability of referral guidelines, training of human resources, involvement of community groups and community-based health education were enablers to improve DRSS by provider side (S5 Table 3).

#### HIC

The main incentives for screening uptake in HICs were awareness of eye care and possibility of treating DR, attending DM education classes, discussion of DM complications with a health care professional, having health insurance with eye care services coverage, higher level of education and being obliged to attend for screening. Adherence to the best practice guidelines by the consumers and having eyes examined by primary care physicians were facilitators.

In HIC availability of educational interventions, DM education programs, adherence to guidelines, targeted screening of high risk groups, reinforcing the importance of eye examination by health care providers, constant screening location, personalised strategies (phone calls or door to door visits), on-line patient access booking system, recall system, showing fundus photograph and teaching patients, ability to change appointments were incentives for uptake of DRSS (S5 Table 4).

#### Quantitative Data Synthesis

The data extracted for quantitative synthesis are available as supporting information S6 Table file.

#### Knowledge and Awareness

The main barriers identified in this systemic review were factors associated with clients. The most consistent barrier across most of the studies was knowledge regarding DR. One study mentioned that knowledge (mean knowledge score 4.7 among those who had examination vs 3.6 without examination (p<0.001) and awareness about DR associated with seeking screening services (OR 1.52, 95%CI 1.1-2.1, p=0.01).(34) This concept was further emphasised in an RCT conducted to evaluate the effectiveness of a health educational intervention and in the intervention arm participants had higher odds of eye examination status (OR 4.3, 95% CI 2.4-7.8). (47) A study from China showed that having a higher DR knowledge score was a potential predictor for ever had an eye examination. (OR 1.31, 95% CI 1.1-1.5, P<0.001).(44)

In a study conducted to enhance the compliance with DRS recommendations, it was seen that when PwDM were given educational material and a notification there was a significant difference in screening uptake (OR 1.4, McNemars x2=102.7; P < 0.0001).(94) A study conducted in Tanzania showed that those who had knowledge on damages to eye due to DM had higher odds of undergoing dilated fundus examination in the past year (OR 19.7, 95% CI 7.0-55.2).(33)

Even the knowledge on DM alone was associated with uptake of DRS. A study mentioned that less practical knowledge about DM (OR 1.5, 95% CI 1.2-2.1) was a factor associated with non-adherence.(84) On the other hand another study mentioned that knowledge on effects of DR on vision was an incentive for uptake of DRS (OR 3.3, 95% CI 2.0-5.5).(88) It was seen that awareness on possibility of treating DR was an incentive for attending screening (OR 1.6, 95% CI 0.9-3.0).(88)

#### Factors associated with awareness

Most of these studies analysed the various patient characteristics and disease factors associated with awareness. Lack of awareness was associated with older age (OR 10.4, p=0.03), poorly controlled HbA1c (OR 4.9, p<0.001) and male gender (OR 1.2, 95% CI 0.7-1.8, p=0.47).(58)

Huang O.S et al in the year 2013 showed that unawareness was associated with lower education (primary or less, adjusted OR 1.9, 95% CI 1.4-2.5, p<0.0001), lower income (Singapore $ <2000, adjusted OR 1.7, 95% CI 1.2-2.5, p=0.003) and poorer literacy (unable to write - adjusted OR 1.4, 95% CI 1.0-2.0, p=0.03).(95)

Thapa, R. et al mentioned that literate patients are more likely to have awareness on DR (OR 2.7, 95%CI 1.3-5.6, p=0.006).(32) One study showed that higher awareness on DR was seen among the more educated people (OR 1.8, 95% CI 0. 9-3.4, p=0.0000).(45) Those who have a history of prior fundus evaluation elsewhere (other than retinal clinics in this study) had higher odds of having awareness on DR (OR 11.9, 95% CI 5.7-25.2) p<0.001).(32)

Thus, it can be assumed that clients’ knowledge and the awareness are main barriers to access. Further health educational intervention may improve the uptake of services. However, the uptake of DRSS can be affected by various socio-economic factors.

#### Attitude

It was also reported that PwDM themselves may not have the judgemental ability over seeking care and recommendation by the provider would improve access (OR 341, 95%CI 164-715, proportion of attendees 99.4% vs non-attendees 34.5%).(88) In health seeking behaviour, those who thought eye examinations were needed every 6 months (OR 1.2, 95% CI 1.1-1.4) and those who worry much regarding their vision (following telephone call intervention, OR 3.47, 95% CI 1.8-6.8) showed higher odds of uptake DRS.(51)(96) A study showed that perception of a PwDM should have eye examination every 12 months (OR 2.62, 95% CI 1.7-4.1, p<0.0001) associated with previous dilated eye examination.(68) Another study showed that fear among patients on impaired vision was an incentive for DR screening (OR 1.9, 95% CI 1.5-2.5).(88)

#### Secondary outcomes of quantitative data - Factors associated with uptake of screening / adherence / regular follow up

##### Service user costs

One major factor associated with undergoing DRS was having an insurance scheme. This was observed mainly in the paid systems. Having an insurance coverage (OR 2.2, 95% CI 1.2-4.3 p=0.02) was associated with the compliance to annual eye examination (85), increased eye screening (OR 3.2, 95% CI 2.2-4.7, p=0.00)(59) and higher chance of undergoing screening (adjusted OR 1.7, 95% CI 1.4-2.2)(75). It is shown that those who have vision loss (blindness) are 100% willing to pay for the services (mean amount willing to pay-No DR - Taiwan dollars - NTD 468.9 ± 327.7 vs Blindness NTD 822.2 ± 192.2, p=0.0005).(86)

One RCT showed that PwDM are less likely to undergo DRS when a co-payment is applied compared to the free services (OR 0.6, 95% CI 0.5-0.7).(65) Those who had no health insurance (OR 2.5, 95% CI 1.7-3.7) were non-compliant for screening.(77) Sheppler, C.R. et al mentioned that those who had an insurance coverage complied with annual eye examination (OR 2.2, 95% CI 1.1-4.3, p =0.02).(85) Lian, J.X. et al showed that being in the pay groups was negatively associated with uptake of screening (OR 0.6, 95% CI 0.5-0.7).(65) Study done by Moss, S.E. et al showed that having a health insurance with eye examination covered (OR 3.3, 95% CI 2.2-5.1, p<0.0001) associated with previous dilated eye examination.(68)

#### Family income

Higher family income was associated with having had a dilated eye examination (US $ >50,000 vs US $ <40,000, OR 1.9, 95% CI 1.3-2.9)(49) (US $ >35,000, OR 1.3, 95%CI 0.8-2.2)(93) The study done by Paskin-Hall, A. et al showed that those who have a higher income ($35,000-$49,000 adjusted OR 1.3, 95% CI 1.1-1.5) had higher odds of undergoing DRS.(75) Another study done in Korea by Rim, T.H. et al showed that those who had highest monthly income quintile (OR 1.4, 95% CI 1.1-1.8, p<0.01) had higher odds of undergoing screening.(79)

#### Gender

The odds of having had a dilated funduscopy in the past year was high among women (OR 1.2, 95% CI 0.9-1.5)(49) and past eye care use decreased by being male (OR 0.5, 95% CI 0.3-0.8, p<0.01)(48). However a study done in UK showed males had higher odds of attending screening following invitation (OR 1.4, 95% CI 1.1-1.7).(54) Therefore, role of gender could be context specific with regard to uptake of DRS.

#### Age

The increasing age (persons with 70 years of age) had twice the likelihood of having undergone dilated fundoscopy compared with those <40 years of age (OR 1.9, 95% CI 1.5-2.6)(49). Most of the studies showed that older PwDM had higher odds of undergoing screening (age >65 years, OR 2.6, 95% CI 1.6-4.1)(93), (OR 1.02, p<0.001).(69)

It was observed that past year eye care use decreased in those who are younger (age 20-39 yrs, OR 0.1, 95% CI 0.01-0.70 p<0.05).(48) Similarly a study done in UK showed that younger age associated with non-attendance (18-34 years, adjusted OR 1.4, 95% CI 1.1-1.7), (35-44 years, OR 1.4, 95% CI 1.2-1.7).(54)

#### Level of education

Most of the studies mentioned the association between level of education and DRS uptake. PwDM with more than high school education vs less than ninth grade education (OR 1.5, 95% CI 1.0-2.1) was associated with higher likelihood of having a dilated eye examination.(49) The reasons for non-adherence mentioned in another study was education less than high school (OR 1.5, 95% CI 1.1-2.1) (77). It was observed that past eye care use decrease by less number of years in educational attainment (<10 years, OR 0.4, 95% CI 0.2-0.9, p<0.05).(48)

In addition education up to high school or more was a predictor of knowledge that uncontrolled diabetes could cause eye disease (OR 2.4, 95%CI 1.5-4.0, P<0.05).(70) Xiong, Y. et al showed that higher awareness on DR was seen among the more educated people. (OR 1.8, 95% CI 0. 98-3.44, p=0.0000).(45)

#### Disease factors associated with uptake of screening

Higher level of glycosylated haemoglobin was associated with non-compliance of screening (>9%, OR 1.7, 95% CI 1.1-2.6).(77)

#### Diabetes / Eye care education

One study mentioned that those without DM education (OR 0.4, 95%CI 0.2-0.6) are less likely to undergo screening.(41) Hwang, J. et al in Canada showed that increased eye screening was associated with discussion of DM complications with health professional (OR 2.0, 95% CI 1.3-3.2, p=0.00).(59) Persons having attended a DM education class (OR 1.5, 95% CI 1.2-1.9) were highly likely of having a dilated eye examination.(49)

A study done in USA showed that eye care education (OR 1.6, 95% CI 1.2-2.1) was associated with receipt of dilated eye examination.(93) No formal DM education (OR 1.3, 95% CI 1.1-1.6) and less practical knowledge on DM (OR 1.6, 95% CI 1.2-2.1) were associated with non-adherence. (84) Those who had attended a DM education class had higher odds of having a dilated eye examination in the past one year (OR 1.5, 95% CI 1.2-1.9).(49) It is shown that when there is no education on DM, patients are less likely visit an ophthalmologist on regular basis (OR 0.39, 95% CI 0.24-0.65).(41)

#### Personnel conducted last eye examination

The reasons for non-adherence to DRS described include personnel conducting last eye examination. It is shown that non-adherence was high when last examination had been conducted by a non-ophthalmologist personnel (OR 4.3, 95% CI 2.3-6.2).(84)

#### Duration of diabetes

The duration of DM and regularity of the clinic visits were predictors of having DR. Those who have DM for less duration (<5 years, OR 0.04, 95% CI 0.01-0.10) were less likely to be not having DR. (36) One study mentioned that those PwDM with <5 years (OR 0.4, 95% CI 0.3-0.8) of duration after diagnosis are less likely to undergo screening.(41)

Hwang, J. et al in Canada showed that duration of DM longer than 10 years (OR 1.5, 95% CI 1.04-2.25, p=0.03)] was associated with increased eye screening.(59) Similarly Saadine, J.B. et al mentioned that when the DM disease duration was longer (>15 years, OR 1.9, 95%CI 1.4-2.6, p<0.0001) those were highly likely to attend for follow ups.(80)

Factors positively correlated with eye care use were time since diagnosis of DM (20 years, OR 2.7, 95% CI 1.2-5.9, p=0.041).(48) In contrast to other studies it showed that when the duration of DM goes up less likely to attend screening (5 to 9 years, OR 1.9, 95% CI 1.6-2.2), (>20 years, OR 3.4, 95% CI 2.7-4.2).(54)

#### Type of diabetes treatment

Factors positively correlated with eye care use was having had treatment with oral antidiabetics+insulin (OR 2.8, OR 1.1-7.4, p=0.161).(48)

#### Regularity of clinic visits

It was observed that past year eye care use decrease those who are younger age (age 20-39 yrs, OR 0.09, 95% CI 0.01-0.70, p<0.05), never married (OR 0.14, 95% CI 0.03-0.76, p<0.05) and having less number of years in educational attainment (<10 years, OR 0.37, 95% CI 0.16-0.88, p<0.05).(48) Further noncompliance was associated with those who had no routine physical examination > 1 year ago (OR 1.8, 95% CI 1.3-2.5).(77)

Mukamel DB et al showed patients who visit their PCPs more often (OR 1.3, 0.001<p<0.01) had high probability of screening in the past 12 month period.(69)

#### Marital status

It was observed that past eye care use decrease in those who were never married (OR 0.14, 95% CI 0.03-0.76, p<0.05).(48)

#### Unemployment

Unemployment was inversely associated with eye care use (OR 0.5, 95% CI 0.2-1.1, p=0.091).(48)

#### Alcohol intake

Heavy alcohol consumption was inversely associated with eye care use (OR 0.3, 95% CI 0.1-0.7, p=0.003).(48)

#### Having other complications of diabetes

Factors inversely associated with eye care use was having diabetic foot (OR 0.4, 95% CI 0.2-0.9, p=0.35).(48)

#### Physician recommendation

Physician recommendation is a predictor of having regular eye examination as mentioned in one study done in Ireland (OR 1.3, 95% CI 1.1-1.6).(51) Van-EijK K.N.D. et al showed that recommendation by the care provider was a strong incentive for undergoing DRS (OR 341, 95% CI 164-715).(88) Similar association has been mentioned by the Wang D et al. (OR 2.2, 95% CI 1.5-3.3, P<0.001).(44)

#### Having other eye diseases and visual impairment

Those who have other eye diseases (OR 1.2, 95% CI 1.1-1.6) and those who think that eye examinations are needed every 6 months (OR 1.2, 95% CI 1.1-1.4) showed higher odds of uptake DR screening.(51) Study done by Moss SE et al 1995 showed that history of cataract (OR 2.9, 95%CI 1.9-4.4, p<0.0001) associated with previous dilated eye examination.(68) Hwang, J. et al in Canada showed that increased eye screening was associated with having visual impairment (OR 2.6, 95% CI 1.7-3.9, p=0.00).(59)

#### Social deprivation

Even in HIC people lived in deprived areas were failed to attend DRS (OR 2.3, 95% CI 1.9-2.8).(63) One study stated that people living in most deprived areas (OR 1.2, 95% CI 1.2-1.3) were not adhered with screening recommendations.(74) A study done in UK showed the factors associated with non-attendance following invitation to screening in a sample of 31,484 diabetics in a DR screening program. In this study social deprivation (adjusted OR 1.4, 95% CI 1.2-1.6, p<0.001) was associated with non-attendance. (54)

Another study done in Korea by Rim, T.H. et al showed that those who lived in urban areas (OR 1.5, 95% 1.2-1.8, p<0.01) and had highest monthly income quintile (OR 1.4, 95% CI 1.1-1.8, p<0.01) had higher odds of undergoing screening.(79) Scanlon, P.H. et al mentioned that each increasing quintile of socioeconomic deprivation was associated with probability of having been screened for DR was decreased (OR 1.1, 95%CI 1.1-1.2, P<0.001).(81)

#### Risk of development of DR among non-attendees

The relative risk of having DR was high in non-attendees as shown in one study conducted in Yemen. (Relative risk of having DR 1.5, 95% CI 1.2-2.2), (bilateral blindness 4.0, 95% CI 1.4-11.6) (low vision disability 2.4, 95% CI 1.8-3.5)(36).

Saadine, J.B. et al mentioned that when those who have moderate or worse retinopathy (OR 2.2, 95%CI 1.6-2.9, p<0.0001) were highly likely to attend for follow ups.(80)

### Discussion

This is the first systematic literature review which describe the barriers and facilitators to access DRS. This evidence will be useful to identify potential barriers and facilitators to implement DRSP in similar settings. Most of the available reviews describe interventions to promote DRS uptake.(97) In addition, there was another Cochrane review on quality improvement interventions to increase DRS attendance.(98) Though there are potential benefits to PwDM, DRS attendance is at a sub-optimal level in screening programs even in HIC settings.(99) Diabetic retinopathy screening has been shown to be cost effective in terms of sight years preserved.(100) Most parts of the world DRS remained non-systematic.

We conducted a narrative synthesis of observational data to understand the barriers and facilitators operationalised in an eye care system when accessing DRSS. This allowed us to conceptualize obstacles for accessing care and to identify potential strategies to overcome those barriers. The findings from this narrative review will be useful to emphasise the challenges that could be faced by users and providers in a DRSP. This will be helpful to explore the avenues for successful implementation of a DRSP in a country.

This review included 63 articles from diverse settings. We used a comprehensive approach to capture all possible articles on this review question. Inclusion of studies without restricting the study design allowed to derive a wide range of themes. We used narrative synthesis of data due to high heterogeneity among the studies. Further we assumed a provide a wide range of barriers and enablers themes by incorporating both qualitative and descriptive quantitative studies.

In order to maintain the homogeneity among the included studies, we divided the studies according to the income setting. This review identified numerous barriers and facilitators to access DRS in a context. Further we explored whether there were differences in barriers between different income settings. In the HIC most of the barriers to access were related to processes of DRS while in LIC and LMIC they were related to major system factors such as lack of human resources and infrastructure. A majority of the studies were focused on the perspectives of the users when describing the barriers. Almost non of the studies explored the perspectives of the policy makers or program planners. Therefore, this review lacks several aspects of stakeholder perspectives.

Synthesis of existing evidence helped to narrow down barriers in to identify modifiable themes. In general, knowledge appeared as the main modifiable barrier to access from user side. However, in a paid system, low income and financial constrains had been mentioned frequently. In addition, most frequently mentioned (i.e., frequency of a theme appear in a study) barrier by users was asymptomatic nature of DR as shown in harvest plot in S2 Fig 2. Complementary to these outcomes, the most common incentives mentioned in included studies were better knowledge on DR / DRS, higher level of education, presence of symptoms and high level of income as shown in S3 Fig 3.

When considering the frequently cited barriers by providers, deficiencies on educating users on DR / DRS, issues in accessibility in making appointments, long waiting time at eye clinics emerged as main barriers (S4 Fig 4). Enablers for providers were educating the users on regular eye examination and providing better access for PwDM (S5 Fig 5).

#### Limitations

The included studies reflected the barriers in a cross section of a time. All the studies used diagnosed PwDM at institutional level as their study samples. There were no studies that used long term sociological and ethnographic approaches to study barriers to access long term in their natural environment.

Many of the barriers or enablers identified in this review was peculiar to modality of screening in the local context. We used reductionistic approach in this narrative synthesis without further synthesis of new themes. Another aspect is the barriers or enablers were assessed in different health systems which has different socio cultural and economic back grounds. Therefore, we could not assess the interactions in between each theme we derived. Though we simplified and de-contextualised the barriers themes, generalizability may depend on the context.

One of the limitations of this review is lack of eligible randomised controlled trials on this review question and primary outcomes were described as explained by the authors. Considering the paucity of systematic reviews under this topic, it is difficult to compare and comment our findings.

#### Implications and public health significance of the findings

Diabetic retinopathy screening program implementation involves a high capital expenditure. There will be a high level of financial risk when implementing a program for the first time. By knowing the potential barriers, the risks can be minimised, and access can be improved by implementing interventions to overcome potential barriers.

The outcomes of current review will be useful to identify the modifiable barriers which could be further explored in a local context before implementing costly interventions. Identification of user and provider perspectives together will enable to identify and cater needs of demand side as well as supply side.

The results of this review show that there are modifiable barriers such as knowledge of DRS among the PwDM which could be use in development of health promotional strategies.

This review highlights the gaps in evidence in this topic in LIC and LMIC. Further there was limited evidence on system factors and stakeholders’ perspectives.

### CONCLUSION

The evidence in this review clearly suggests the barriers and enablers in different income settings. Most consistent barrier across most of the income settings was lack of knowledge and awareness on DR and DRS among the users. In providers point of view, lack of skilled human resources and screening infrastructure was the main barrier. Knowing the modifiable barriers in a context may be helpful to improve the DRSS uptake in institutional PwDM.

## SECTION 3

### Supporting information

S1 Fig 1. Schematic diagram of “Social Cognitive Theory”

S2 Fig 2. Harvest plot showing user barriers

S3 Fig 3. Harvest plot showing user incentives

S4 Fig 4. Harvest plot showing provider barriers

S5 Fig 5. Harvests plot showing enablers for providers

S1 Table. PRISMA check list

S2 Table. Search strategy of barriers to access systematic review

S3 Tables. Methodological quality and applicability assessment of the included studies

S4 Tables. Themes tables by country income setting

S5 Table. Quantitative Data Synthesis - Factors associated with DR screening uptake and regular follow up

S6 Table. Participants’ characteristics of included articles

## Acknowledgements

(None)

---

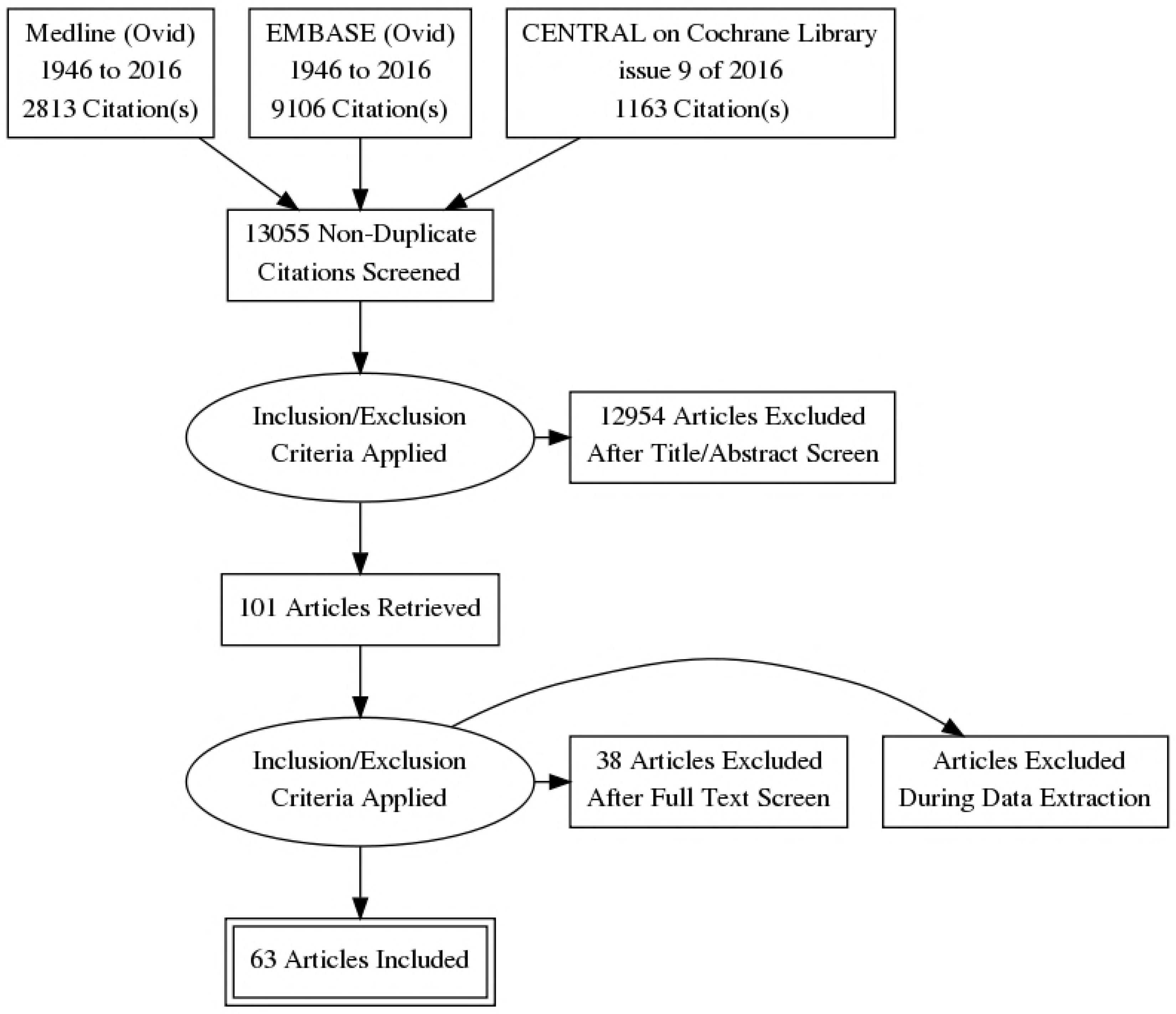

